# Experience-dependent modulation of collective behavior in larval zebrafish

**DOI:** 10.1101/2024.08.02.606403

**Authors:** Roy Harpaz, Morgan Phillips, Ronan Goel, Mark C. Fishman, Florian Engert

**Author notes:** Correspondence and requests for materials should be addressed to R.H.

## Abstract

Complex group behavior can emerge from simple inter-individual interactions. Commonly, these interactions are considered static and hardwired and little is known about how experience and learning affect collective group behavior. Young larvae use well described visuomotor transformations to guide interindividual interactions and collective group structure. Here, we use naturalistic and virtual-reality (VR) experiments to impose persistent changes in population density and measure their effects on future visually evoked turning behavior and the resulting changes in group structure. We find that neighbor distances decrease after exposure to higher population densities, and increase after the experience of lower densities. These adaptations develop slowly and gradually, over tens of minutes and remain stable over many hours. Mechanistically, we find that larvae estimate their current group density by tracking the frequency of neighbor-evoked looming events on the retina and couple the strength of their future interactions to that estimate. A time-varying state-space model that modulates agents’ social interactions based on their previous visual-social experiences, accurately describes our behavioral observations and predicts novel aspects of behavior. These findings provide concrete evidence that inter-individual interactions are not static, but rather continuously evolve based on past experience and current environmental demands. The underlying neurobiological mechanisms of experience dependent modulation can now be explored in this small and transparent model organism.

## Introduction

Collective behavior is crucial for animal survival, and can emerge from simple local interactions among individuals (1–8). Understanding the exact nature of these local interactions, or the behavioral ‘rules’ that animals use to respond to their group mates has been the focus of extensive experimental and theoretical studies (1–14).

Importantly, moving animal groups, such as schooling fish or flocking birds, can exhibit a diversity of stable collective states such as schooling (directed group motion), swarming (disordered motion) and milling (circular motion around a center point) and the ability to dynamically switch between states. Theoretical and experimental studies indicate that such diversity of collective states can emerge from local interactions among individuals that are commonly considered static and hardwired. Specifically, simulated groups of moving agents exhibit different emergent collective states depending on the initial and boundary conditions of the system (2, 15–18). Experimental evidence in schooling fish support these theoretical findings, indicating that groups can transition between multiple stable behavioral regimes, or states, while implementing static interactions among individuals (19–24).

However, assuming inter-individual interaction rules to be static and hardwired ignores the fact that all living systems require plasticity and feedback regulation to maintain homeostasis. In particular, social interactions need to be flexible to allow individuals in groups to respond to evolving social and environmental demands. Yet, little is known about if and how animals can adaptively modulate their inter-individual interactions and the resulting collective behaviors, based on previous experiences, internal states and behavioral goals.

One key macroscopic feature of groups that animals may need to adaptively control is population density (25). Group density is a prominent factor that affects key animal behaviors, such as foraging (26–28), mating (29) and predator avoidance (27, 28, 30), and affects parasite and disease transmission (31). The need for density sensing and control seems to be a general feature for all taxa, from bacteria that use quorum sensing to assess local population density (32, 33) to humans responding to pedestrian flow (34). To achieve such control, animals must be able to continuously sense or estimate the *density* of neighbors in their current environment, overcome local fluctuations and noise in their assessment and to adaptively modulate their behaviors in response to these estimates (1, 35–38). The mechanisms that allow fish to sense, internalize, and change behaviors in response to different population densities are not known.

Larval zebrafish is an emerging animal model in behavioral neuroscience, and recently, the inter-individual interactions underlying collective behavior in larval, juvenile and adult zebrafish were described (10, 12, 13, 38–40). It was shown that at younger ages, up to 7dpf (days post fertilization), these fish exhibit primarily repulsive inter-individual forces which lead to overdispersed group structures. Around 14 dpf, attractive forces and aggregation tendencies start to emerge which become robustly expressed at 21 dpf (12, 13, 41). Yet, little is known about the ability of larval zebrafish to adaptively modify their inter-individual interactions based on recent social experiences. In contrast, long term modulation of low level sensorimotor programs and algorithms was shown to affect individual behaviors such as hunting (42) and defensive strategies (43, 44) already at the larval stage. Therefore, the simple social interactions exhibited at early developmental stages together with the unique ability to study the nervous system at the whole-brain single-cell resolution at these times, offers a unique opportunity to tap into the behavioral and neural mechanisms of experience dependent modulation of collective behavior.

Here, we use naturalistic and virtual-reality (VR) experiments and explore how young larval zebrafish - at 7 dpf - modulate their repulsive interactions in response to persistent changes in population density. We infer the specific visual cues that fish use to estimate their current group density and we precisely track experience-dependent modulation of inter-individual avoidance behavior in response to these estimates. We show that modulation is a slow and stable process that develops within tens of minutes and lasts for many hours. Finally, we present a time-varying computational model that implements experience dependent gain changes in the sensorimotor transformations that connect the visual inputs generated by conspecifics into concrete swim and turn behaviors. This model accurately accounts for the experience-dependent modulation of collective behavior in groups of larval zebrafish, and reproduces our behavioral findings. Our results demonstrate how even at a very early age, when collective social behavior is just beginning to emerge, larval zebrafish are already able to adjust their social interactions and the resulting collective behavior to adaptively cope with changing social conditions.

## Results

### Previous social experience of larvae modulates their collective behavior

To study how recent social experiences of young larvae modulate their collective behavior, we divided 7 dpf (days post fertilization) larvae into groups of either 5 (low density) or 20 (high density) animals, allowed each group to swim together for 4 hours in opaque circular arenas (diameter = 6.5cm) and measured their collective swimming behavior (Fig. 1a-b, Methods). We then redistributed fish into groups of the opposite density and measured their collective swimming behavior again (Fig.1a-b)(Movies 1). This procedure allowed us to compare fish swimming in similar group densities that differ only in their history of social experiences (Fig. 1a, compare dotted blue to solid blue and dotted orange to solid orange).

As previously shown (12, 41), 7 dpf larvae tend to swim in overdispersed groups in both high and low densities, with average nearest neighbor distances that are higher than those of shuffled controls (NN_5_=1.41±0.11 [mean±SD], NN_5_^Shuffled^=1.32, NN_20_=0.65±0.03, NN_20_^Shuffled^=0.6 dashed horizontal lines, p_5_=3. 4 · 10^−6^, p_20_=5. 6 · 10^−6^, Wilcoxon sign rank test)(Fig. 1c-d). When comparing the effects of previously experienced social density on the current behavior of fish we found that group structure was modulated by past social experience: Fish swimming in low density groups that were previously exposed to high density swam closer to their neighbors (NN_5_=1.41±0.11, NN_20→5_=1.34±0.1 [mean±SD], p=0.0007, Wilcoxson rank sum test, Cohen’s d=-0.65), and conversely, fish swimming in high density groups that were pre-exposed to low density swam farther away from one another (NN_20_=0.65±0.03, NN_5→20_=0.67±0.02, p=0.0462, Cohen’s d=0.58)(Fig. 1c-d, Fig. S1a). Importantly, previous social experiences did not consistently change other individual and group properties such as distance to the walls, average swimming speed, average distance traveled in a bout and the bout rate of the fish (Fig. S1b-f).

The change in inter-individual distances due to previous social experience is likely the result of the modulation of inter-individual interactions. Larval zebrafish are known to use retinal occupancy of neighbors to guide their inter-individual interactions (12). Specifically, 7 dpf larvae were found to turn away from the more occupied eye, which was shown to fully explain the over-dispersion phenotype in these young animals (12, 41).

Here, we confirmed these results for larvae swimming in groups of either 5 or 20 animals, and show that fish in our experiments also tend to turn away from the more occupied eye (Fig. 1e). Critically, we found a history dependent modulation of these inter-individual interactions that explains the changes in collective swim behavior described in Figure 1c,d: larvae swimming in groups of 5 animals showed a reduced tendency to turn away from the more occupied eye, after experiencing a higher density (experimental condition 20→5), while larvae swimming in groups of 20 animals showed an increased tendency to turn away, after experiencing a lower density (experimental condition 5→20)(Fig. 1e).

Taken together, these results show that collective behavior depends on the previous social experiences of the larvae, and that fish can either increase or decrease the strength of their social interactions, depending on the specific experienced history.

**Figure 1.**
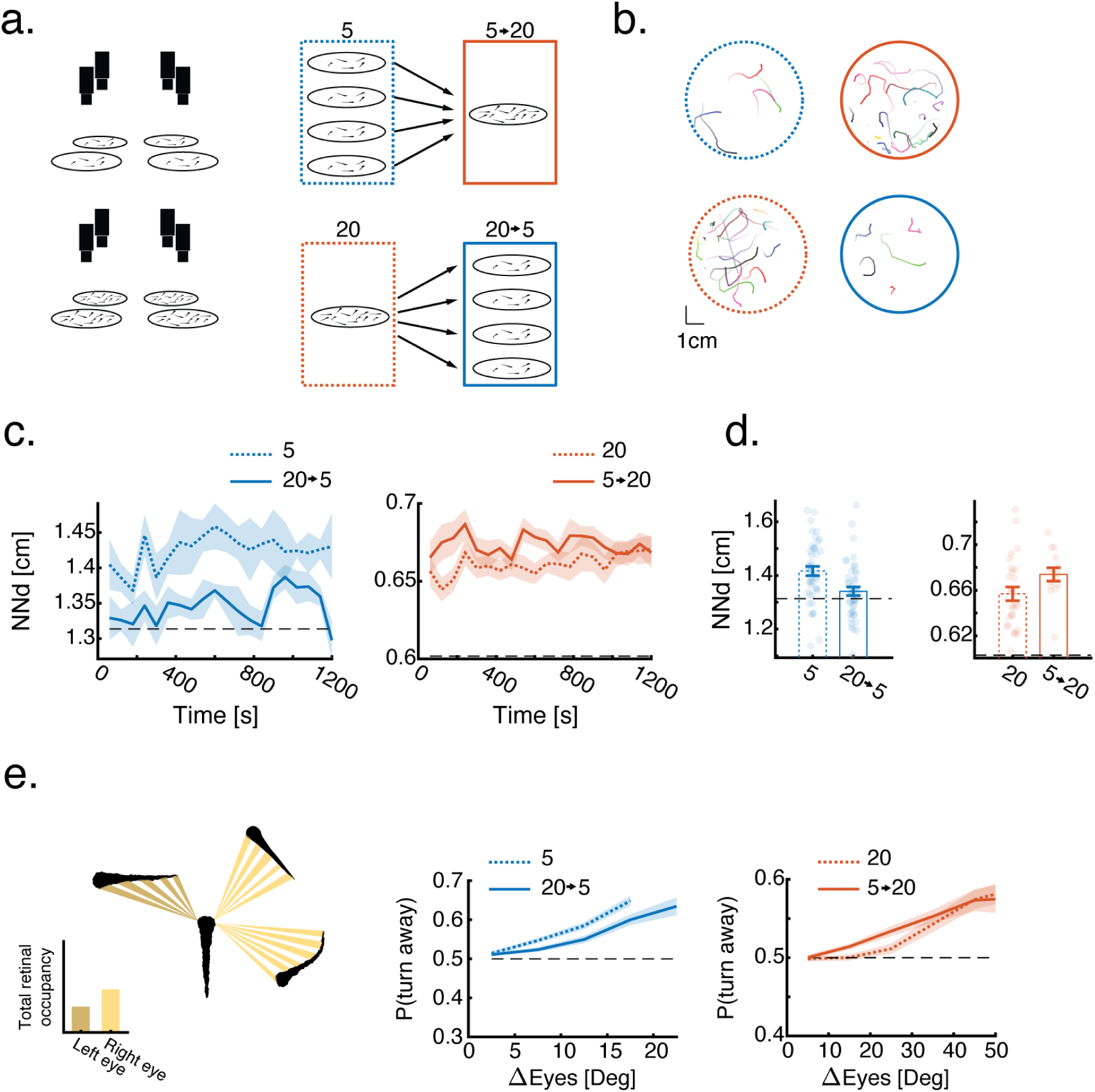
Previous social experience modulates collective swimming behavior. **A.** High throughput experimental design to study experience dependent modulation in groups. Fish behavior is monitored in high and low densities. Groups are then combined or split to study the opposite density. **b.** Example of the joint trajectories of groups of 5 or 20 fish over 20 seconds. Colors represent different fish. **C.** Time dynamics of the average nearest neighbor distance of groups swimming in either low (dashed blue) or high (dashed red) neighbor density, and for groups that were previously exposed to the opposite density (solid lines). Individual group data are averaged in 60s blocks. Shaded areas are SEM. **d.** Average nearest neighbor distance of groups (N_5_=44, N_20→5_=45, N_20_=27, N_5→20_=14). Dots represent individual groups, error bars are mean±SEM. Black dashed lines represent average values of shuffled controls (Methods). Previous experience with an opposite social density modulates group structure. **e.** Probability to turn away from the more occupied eye as a function of difference in retinal occupancy (higher occupied eye - lower occupied eye). Lines represent mean turning probability calculated as the fraction of turns away from the more occupied side out of all turns in 5° (groups of 5) or 10° (groups of 20) bins. Shaded areas are SEM. Previous social experience modulates fish responses to retinal occupancy.

### Interactions with virtual neighbors elicit similar modulations of collective behavior

To precisely characterize the history-dependent modulation of larval collective behavior and to infer the mechanisms at the basis of this modulation we utilized a high throughput virtual-reality (VR) assay that allowed us to study how larvae modulate their behavior when exposed to different spatial and temporal retinal occupancy patterns (Fig. 2a, Movie 2, Methods). To this end we projected dots that mimicked the size and motion of real groups of fish, and measured the behavior of a single fish interacting with these virtual neighbors (Fig. 2a-b, S2a, Methods). This virtual-reality assay allowed us to easily transition between neighbor densities (Fig. 2a, right) without the need to physically disturb the fish or move them between groups, and also to fully control the physical and dynamic properties of the simulated virtual neighbors (see Fig. 3 and below).

To test the effects of previous social experiences in virtual-reality on inter-individual distances, we first exposed fish to either a high or low density of virtual neighbors (4 or 19 virtual neighbors, Methods) (Fig. S2a). We then switched the virtual neighbor densities (from low to high or vice versa) and recorded the responses of the fish before and after density switches (Fig. 2b).

We found that similar to experiments with real groups of larvae, fish swimming with 4 virtual neighbors that were previously exposed to 19 virtual neighbors swam closer to their neighbors compared to fish that did not experience a change in density (NN_5_=1.59±0.17, NN_20→5_=1.45± 0.09 [mean±SD], p=0.007 Wilcoxson’s rank sum test, Cohen’s d=-1.03)(Fig. 2c, S2b). In contrast, fish that swam with 19 virtual neighbors and were previously exposed to 4 virtual neighbors swam farther away from their neighbors (p=0.0546, NN_20_=0.6±0.04, NN_5→20_=0.63± 0.04 [mean±SD], Wilcoxson’s rank sum test, Cohen’s d=0.75)(Fig. 2c, S2b). Similar to the group swimming experiments, changes in virtual densities did not consistently affect other swim properties such as larvaes’ distance to the walls, swim bout rates and distance traveled in each bout (Fig. S2d-f).

Here again, we could trace the change in inter-individual distances to the strength of responses to difference in retinal occupancy between the eyes: fish that were previously exposed to high density of virtual neighbors, exhibited a reduction in tendency to turn away from the more occupied eye, while fish that previously experienced a lower density of virtual neighbors showed an increase in the probability to turn away from the more occupied eye (Fig. 2d).

These results confirm that visual social interactions control collective behavior of 7 dpf larvae and also that history-dependent modulation of social interaction strength can be reliably induced in a VR setting.

### History dependent modulation of collective behavior is slow, stable and reversible

Next, we analyzed the quantitative dynamics of experience dependent modulation and the stability of behavioral changes. We focused on the modulation caused by previous exposure to high densities, as it resulted in a more robust change in group behavior (Fig. 1c-d and Fig. 2c). To measure the temporal dynamics of collective behavior modulation, we compared group behavior at low neighbor density (4 virtual neighbors) before and after exposing the fish to varying times (5 to 40 minutes) of high neighbor density (19 virtual neighbors)(Fig. 2e, Methods). We found that longer exposure times to high neighbor density caused a stronger decrease in the distances that fish kept from their virtual neighbors (Fig. 2e). Surprisingly, the decrease in inter-individual distances due to exposure to high neighbor density (19 virtual neighbors for 20 min), remained stable for more than 2 hours after switching back to a low neighbor density (Fig. 2f).

To explore ways to reverse these behavioral modulations, we tested whether subsequent exposure to the lowest possible density (0 neighbors), can negate the effects of exposure to high neighbor density. Indeed, we found that increasing times of exposure to 0 neighbors can reverse this effect and bring inter-individual distances among fish back to baseline in approximately 50 minutes (Fig. 2g, Fig. S2g-h).

These results indicate that exposure to high density results in a slow cumulative change in the collective behavior of the fish, characterized by closer proximity of neighbors, a phenomenon that remains stable over many hours. This process can be reversed by removing all virtual social stimuli, which causes the inter-individual interactions and distances between fish to slowly decay back to their baseline measured values.

We note that the behavioral adaptations observed in VR must rely exclusively on visual experiences. In order to isolate the precise visual features of neighbors that trigger this modulation we independently tested the ability of specific visual features, such as the object’s density, their retinal occupancy and their motion statistics, to elicit behavioral modulation.

**Figure 2.**
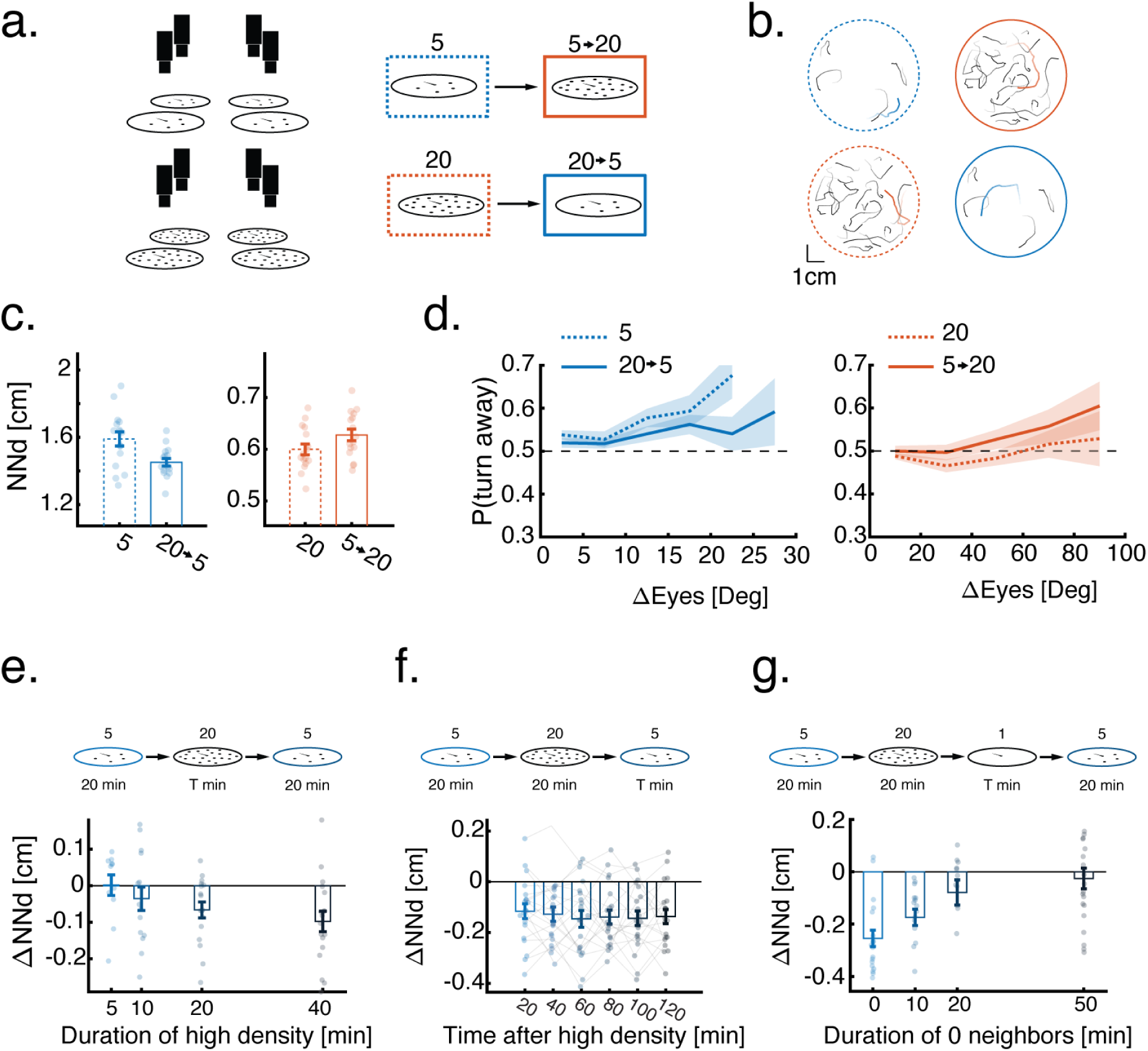
Dynamics of experience dependent modulation. **a.** High throughput VR assay to study experience dependent modulation of collective behavior. Single fish are exposed to virtual neighbors moving according to real, pre-recorded trajectories. The density of virtual neighbors is switched mid experiment. **b.** Example trajectories (over 20s) of single real fish (red/blue) and their virtual neighbors (black) while fish are exposed to either low (blue) or high (red) virtual neighbor densities. **c.** Average nearest neighbor distance of real fish from their virtual neighbors. Dots represent individual fish, error bars are mean±SEM (N=16 for all conditions). Previous experience with an opposite virtual social density modulates nearest neighbor distances. **d.** Probability to turn away from the more occupied eye as a function of difference in retinal occupancy (higher occupied eye - lower occupied eye). Lines represent mean turning probability calculated as the fraction of turns away from the more occupied eye out of all turns in 5° (4 virtual neighbors) or 10° (19 virtual neighbors) bins. Shaded areas are SEM. Previous social experience modulates fish responses to virtual retinal occupancy. **e.** Average change (after-before) in nearest neighbor distances of fish swimming in a low neighbor density after exposure to increasing times of high neighbor density (N_t=5_=11, N_t=10_=15, N_t=20_=17, N_t=40_=18). **f.** Average change (after-before) in nearest neighbors distances of fish swimming in a low neighbor density after exposure to a high density of virtual neighbors (20 min). Bars represent averages over fish in blocks of consecutive 20 minutes after exposure to high density. Lines connect individual fish (N=20). **g.** Average change (after-before) in nearest neighbor distance of fish swimming in a low neighbor density after exposure to 20 min of high virtual neighbor density followed by increasing times of 0 neighbors (N_t=0_=19, N_t=10_=16, N_t=20_=13, N_t=50_=21). In panels (**e-g**) top panels represent experiment conditions, single dots are individual fish and error bars represent mean±SEM.

### Persistent changes in retinal occupancy drive modulation of preferred neighbor proximity

To test which statistics of the visual scene are used by larvae to modulate their turning responses, we varied the size and motion statistics of virtual neighbors independently from their density, and measured their ability to elicit modulation of preferred nearest neighbor distances (Fig. S3a-b). We found that increasing either the size or the speed of virtual neighbors, while keeping their numbers constant, also caused fish to reduce their tendency to turn away from neighbors, and consequently swim closer to each other (Fig. S3a-b). We did not observe any change in group structure or inter-individual distances when replacing virtual neighbors with a whole-field black background as a stimulus, indicating that the modulation is not a simple adaptation to global luminance (Fig. S3c)(45). In addition, presenting fish with high density of stationary neighbors also failed to elicit modulation of preferred neighbor proximity (Fig. 3a, Movie 3).

These results indicate that changes in the perceived motion energy (translational, looming, or both) of neighbors on the retina are at the basis of collective behavior modulation. We therefore sought to determine the specific contributions of each of those components. To that end, we presented fish with two sets of stimuli each isolating either looming or translational motion: a high density of virtual neighbors (19 neighbors) that are stationary but randomly increase and decrease in size (looming, not moving), or a high density of virtual neighbors moving on concentric circles around the focal fish (moving, not looming)(Fig. 3a, Movie 3, Methods).

We found that only changes in looming energy caused a reduction in inter-individual distances, while changes in rotational motion energy were insufficient to induce any modulation (Δ NN_stationary_=0.03±0.13, ΔNN_Translational_=-0.025±0.17, ΔNN_Looming_=-0.09±0.15 [mean±SD]; p_Stationary_=0.394, p_Translational_=0.440, p_Looming_=0.011, Wilcoxon signed rank test)(Fig. 3a, Movie 3). The reduction of inter-individual distances was not associated with consistent changes in other individual swim properties such as the path traveled in a bout or the bout rates of these fish (Fig. S3e-f).

To further corroborate these findings, we directly manipulated translational and looming motion using a single dot stimulus, mimicking a neighboring fish swimming in the vicinity of the focal fish, and tested its ability to elicit modulation of inter-individual interactions (Fig. 3b, Movie 4, Methods)(12). The single dot stimulus moves in bouts ’rotationally’ around the fish from ±75° to ±30° for 5 seconds (with fish heading taken as 0°). We either kept the size and distance of the dot constant to elicit only translational motion on the retina, or slowly changed the size of the dot or its distance to the fish, to add a looming component.

We find that all stimulus sets elicit robust turning responses (Fig. 3b, bottom, Block 1), as shown previously (12) and similar to the case of multiple virtual neighbors (Fig. 2d).

However, the addition of looming, significantly reduced future response probabilities to the single dot stimulus, whereas fish kept responding robustly to a moving dot stimulus with no looming component (Fig. 3b, bottom, Blocks 4-5).

We next tested whether changes in response probability to the single dot stimulus also translate into changes in nearest neighbor distances in a group context. We found that after training with single dot looming stimuli, fish swam closer to their virtual neighbors, while there was no apparent change in neighbor distances after exposure to a non looming stimulus (ΔNN_moving_=-0.01±0.14, ΔNN_moving closer_=-0.05±0.12, ΔNN_moving and growing_=-0.06±0.14 [mean±SD], p_moving_=0.19, p_moving closer_=0.03, p_moving and growing_=0.034, Wilcoxson’s signed rank test)(Fig. 3c).

Taken together, these results demonstrate that modulation of collective behavior is caused by repeated looming events as, for example, evoked by the movement of neighbors in the vicinity of the animal.

**Figure 3.**
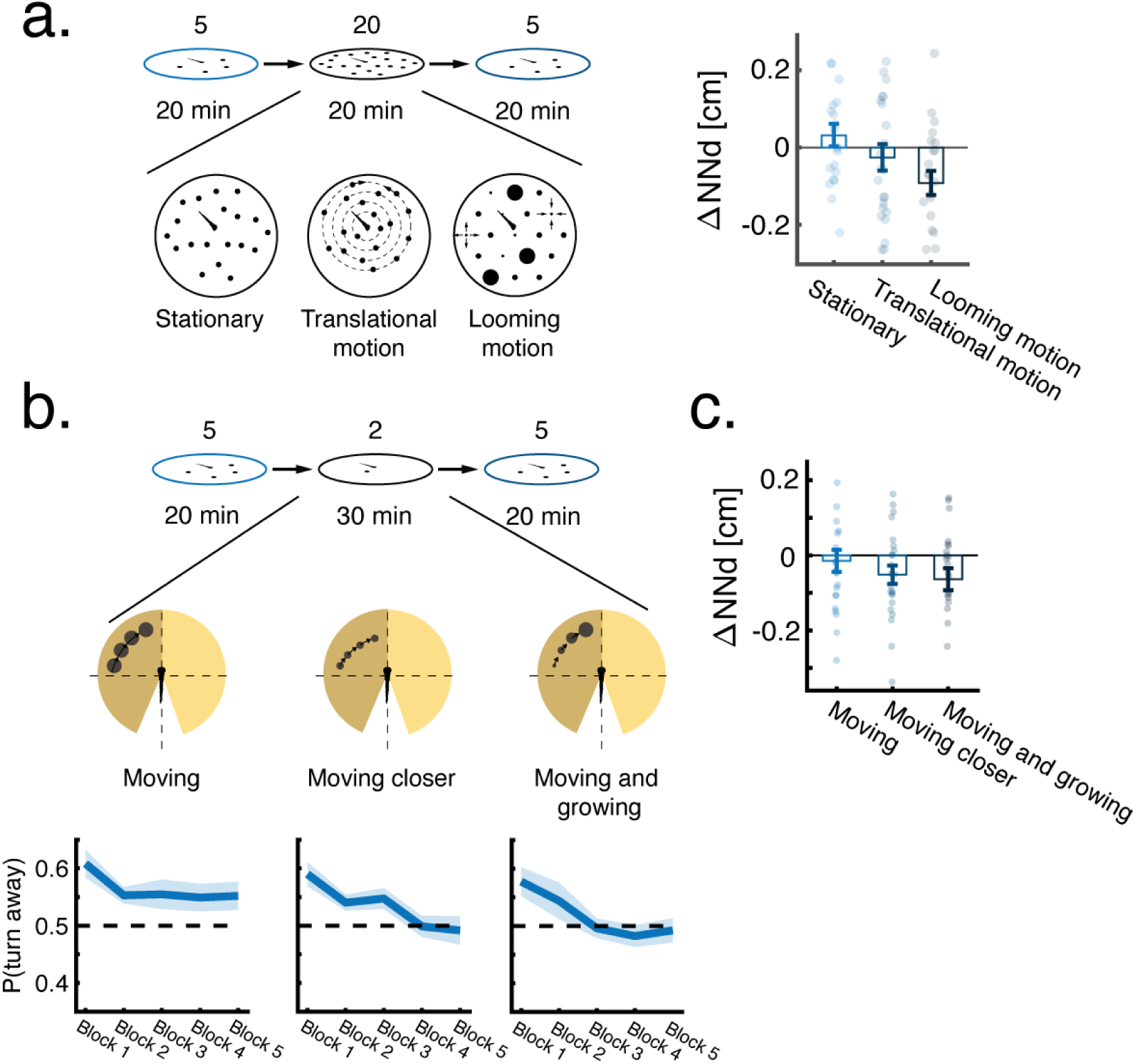
Persistent changes in retinal occupancy drive experience-dependent modulation of collective behavior. **a.** Average change (after-before) in nearest neighbor distances of fish swimming in low density (4 virtual neighbors) after exposure to 20 min of high density (19 virtual neighbors) with different motion statistics (N_stationary_=21, N_rotation_=24, N_looming_=22, Methods). **b. Top:** Experiments testing the ability of a single dot stimulus with different motion statistics to elicit changes in nearest neighbor distance (Methods). **Bottom:** Probability per bout to turn away from a moving dot in the 3 conditions. Bold lines are the mean probability over fish, and shaded areas are SEM. Probability is calculated as the fraction of turns away from the presented side out of all turns, and each block is the average of 60 stimulus presentations. Dotted lines represent no preference in turning direction. **c.** Average change (after-before) in nearest neighbor distance of fish swimming in a low density group after exposure to 30 min of single dot stimuli as shown in **b.** (N_moving_=24, N_moving closer_=24, N_moving and growing_=23). Single dot stimuli with a looming component to their motion cause a subsequent reduction in nearest neighbor distances of fish in a group.

### Modeling experience-dependent modulation of collective behavior

We devised a modeling framework that links visually based inter-individual interactions to the history of experienced visual scenes (Fig. 4a-c). The model has two time varying components - an interaction function 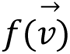 that defines the way a larval zebrafish will respond to the retinal occupancy 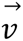 generated by its neighbors, and an unobserved internal state variable *s*(*t*) that represents a cumulated value of previously experienced retinal occupancy changes and modulates the response to visual occupancy 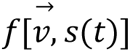.

Previously, we have shown that larval zebrafish base their turning responses on precise retinal occupancy computations: fish spatially integrate retinal occupancy in the vertical dimension at different visual angles and these integrated values elicit directional turning biases. Turn biases are then horizontally averaged within each eye and the resulting values are compared between the two eyes to elicit a turning response (12)(Fig. 4b, Methods). We estimated 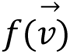 based on these algorithms, in the low density case (4 virtual neighbors) before and after exposure to high density (19 virtual neighbors)(Methods). As expected, the estimated probability to turn away from the more occupied eye is weaker after exposure to high density (Fig. 4b, right).

Next, we linked the transition between stronger and weaker responses, to the history of experienced changes in retinal occupancy by the fish. To that end, we allow agents to update an internal state variable *s*(*t*), by introducing a leaky integrator module that temporally integrates looming events, which we hypothesize to be extracted at the level of each retina and then combined in a downstream retinofugal region (43, 46, 47)(Fig. 4c-d). This module is represented by the linear differential equation:

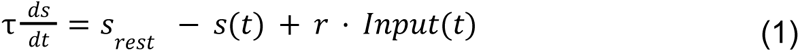

where τ represents the time constant of the process, *Input*(*t*) is the experienced maximal loom at time *t*, *r* is a scaling variable, and *s_rest_* is the value the process will relax to in the absence of any input. Based on the temporal dynamics observed in our experiments (Fig. 2e,g) we equipped this integrator module with a very slow time constant (τ) of ∼6 hours (Methods). We then linearly link the strength of the evoked turning response to this internal state variable, where higher integrated values result in a weaker turning response, and small values of *s*(*t*) in a stronger bias to turn away (Fig. 4c-d, Methods).

We note that, based on the slow time constant, the linear differential equation governing our model (Eq. 1), has a steady state (equilibrium) solution that is only reached after many hours, which implies that the model operates largely in the initial, linear, domain, where sensitivity to changes in the input is high (Methods).

**Figure 4.**
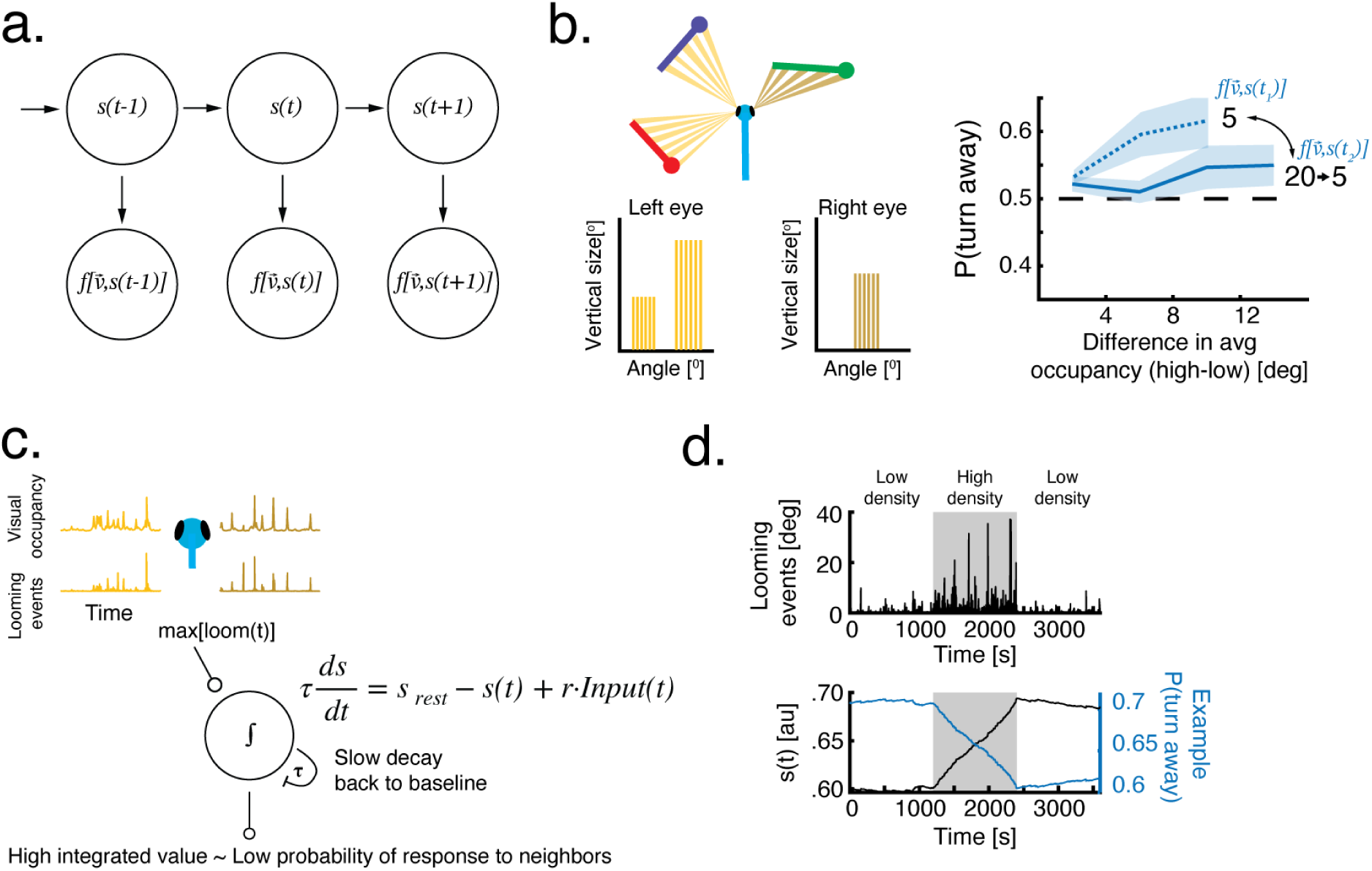
Time varying models of experience-dependent modulation of collective behavior. **a.** A sketch of a time varying model with two components - a hidden state variable *s*(*t*) and an observed response to retinal occupancy of neighbors 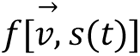 that depends on the hidden state variable. **b. Left:** The response to retinal occupancy 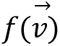 depends on the difference in average retinal occupancy within each eye (12)(Methods). **Right:** Estimating 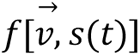 from fish responses to virtual neighbors reveals that fish turn away from the more occupied eye and that responses are modulated due to previous exposure to a different social density (Methods) **c.** A leaky integrator module tracks the values of the hidden state variable *s*(*t*). The input to the integrator is the experienced maximal increases in retinal occupancy (i.e. looming motion on the retina). **d. Top:** Example of experienced looming events when fish are exposed to a low-high-low density experimental design. **Bottom:** The values of the internal state variable *s*(*t*) and the corresponding change in P(turn away) in response to a constant average difference in retinal occupancy. Increasing values of *s*(*t*) cause a reduction in directed turning responses and vice versa.

### The model accurately predicts experience dependent modulation of collective behavior

We tested the ability of the model to explain and predict our behavioral observations (Fig. 1-3). To that end we first quantified the behavior of a single simulated fish (agent) exposed to virtual neighbors of different group sizes, as described for real fish in Fig. 2 (Fig. 5a, Methods). We find that the nearest neighbor distances of an agent interacting with neighbors in low density conditions (4 virtual neighbors, swimming according to the pre-recorded trajectories used in VR experiments) were similar to those exhibited by real fish in VR experiments (Fig. S4a). The transition to high density (19 virtual neighbors) was accompanied by a linear increase in values of the internal state variable (due to the related increase in looming events, Fig. S4a) and a corresponding decrease in the strength of fish turning responses, which resulted in a decrease in nearest neighbor distances (Fig. 5b). This new group structure then remained stable for more than 2 hours of simulation time, in line with the long time constant of the leaky integrator module, which captured the stability of the modulation observed in VR experiments (Fig. 2f).

Increasing the exposure time to high density of virtual neighbors caused a corresponding decrease in the average nearest neighbor distance (Fig. 5c), accurately capturing similar trends observed in VR experiments (Fig. 2e). Furthermore, removing all visual stimulation after exposure to high density, caused the strongest decrease in internal state variable values (Fig. S4a) and an exponential return to baseline nearest neighbor distances in about ∼50 minutes of simulated times (*R_exponential_*^2^ =0.99)(Fig. 5d), which again matched the experimental observations in VR (Fig. 2g).

The model also accounts for more subtle observations in our experiments. Simulated agents exposed to a high density of stationary objects (non-moving neighbors), did not show a decrease in their response to neighbors, or their inter-individual distances, capturing the finding that occupancy changes due to own motion are insufficient to elicit behavioral modulation (Fig. S4b, Fig. 3a). Exposure to even higher neighbor densities (39 virtual neighbors compared to 19 virtual neighbors) caused a stronger decrease in nearest neighbor distances in both experiments and simulation (Fig. S4b, Fig. S3d). Similar to experimental findings, exposure to low density of faster moving neighbors (4 virtual neighbors moving at 4 times the original speed) caused a subsequent decrease in nearest neighbor distances (Fig. S4c, Fig. S3b). In addition, simulated agents exhibited an increase in the distance kept from the walls of the arena after exposure to high neighbor density mimicking the effect observed in real fish (Fig. S4d, Fig. S2h), even though no explicit change to wall interactions was introduced into the model. The model also predicts that priming fish with 0 neighbors before introducing a low neighbor density (4 virtual neighbors) should cause an increase in inter-individual distances. Experiments priming fish with 0 neighbors confirmed this prediction (Fig. S4e).

Finally, we replaced the non-interacting virtual neighbors with groups of fully responsive, interacting agents (Fig. 5e-f, Methods, Movie 6) and compared their behavior to those of real animals swimming together in groups of varying densities (Fig. 1a-d). Here again, agents that were previously exposed to high density groups (20 interacting agents) and were randomly picked out of the cohort and virtually ‘transferred’ to a low density group (Methods), exhibited a decrease in nearest neighbor distances and swam closer to their neighbors (Fig. 5f, S4f). Conversely, simulated agents that were exposed to a low density scenario (5 interacting agents) and were virtually ‘combined’ to create a high density group, swam at higher distances from their neighbors (Fig. 5f, S4f).

Taken together, these results show that simulated agents that are equipped with an internal state variable that integrates and ‘remembers’ the history of experienced retinal occupancy changes, can comprehensively account for most if not all observations of collective behavior modulation in larvae.

**Figure 5.**
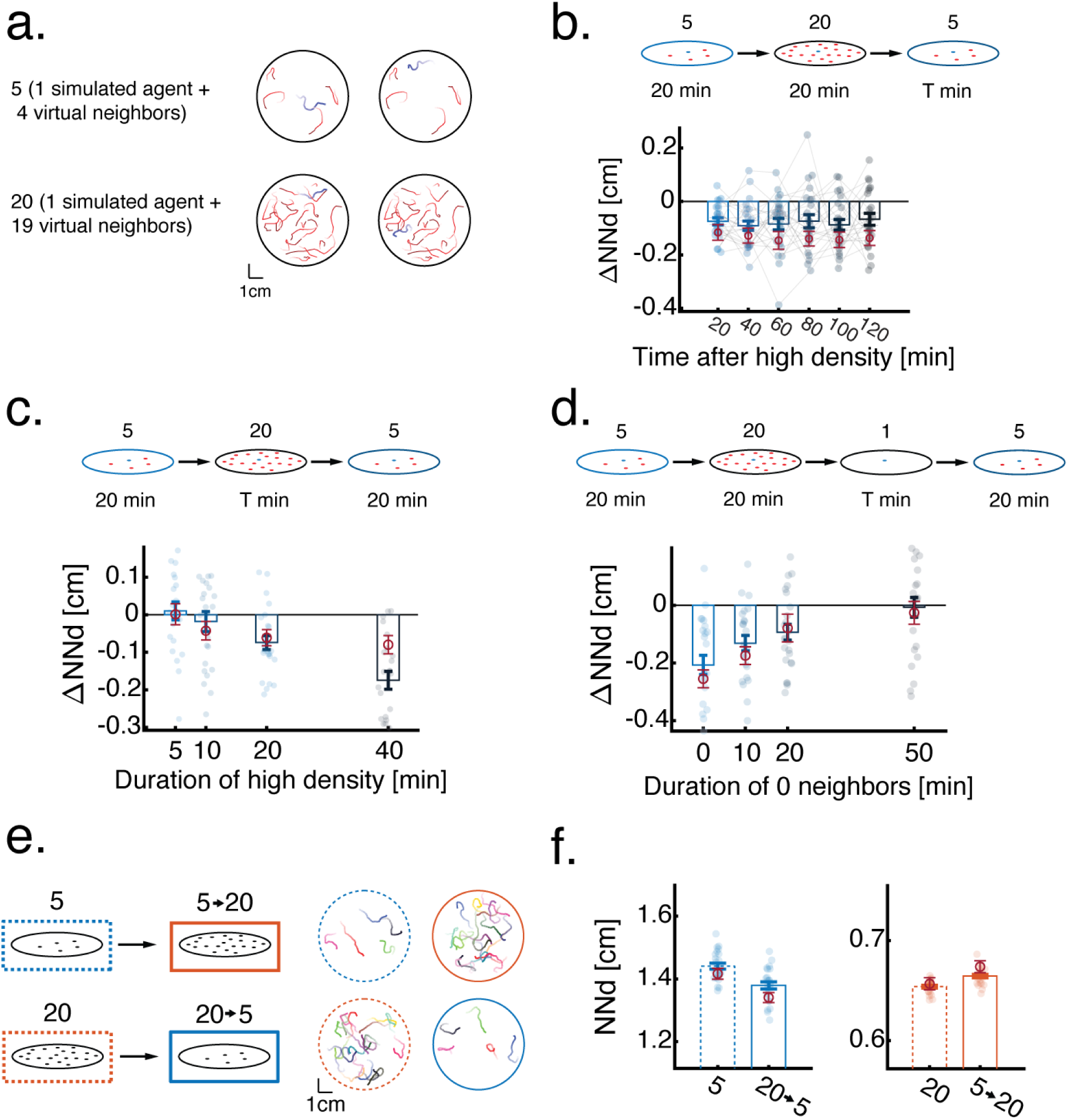
Models accurately predict experience dependent modulation of collective behavior. **a.** Example trajectories (20s) of simulated agents (blue), interacting with ‘virtual neighbors’ (red) based on the model presented in Fig 4. (Methods). **b.** Average change (after-before) in nearest neighbors distances of simulated agents swimming in a low neighbor density after exposure to a high density of virtual neighbors (20 min). Bars represent averages over blocks of consecutive 20 minutes after exposure to high density. Lines connect individual agents. **c.** Average change (after-before) in nearest neighbor distance of simulated agents in a low neighbor density after exposure to increasing times of high ‘virtual neighbor’ density. **d.** Average change (after-before) in nearest neighbor distance of simulated agents in a low ‘virtual neighbor’ density after exposure to 20 min of high ‘virtual neighbor’ density followed by increasing times of 0 neighbors. In panels (**b-d**) top panels represent simulation conditions, single dots are individual agents (N=24), error bars (blue) represent mean±SEM of simulated data. Dark red circles and error bars represent mean±SEM of the real fish data presented in (Fig. 2e-g). **e.** Example trajectories (20s) of groups of simulated agents interacting according to the model presented in Fig. 4 (Methods). Simulated agents are virtually ‘split’ and ‘combined’ to mimic the experimental design used for real groups of fish and presented in Fig. 1a. **f.** Average nearest neighbor distances of groups of simulated agents that swim in low (left - blue) and high (right - orange) group densities, solid lines show simulated groups that were pre-exposed to the opposite density. Dots represent individual groups of agents, error bars are mean±SEM in simulations. Dark red circles and error bars represent mean±SEM of the real fish data presented in (Fig. 1d). N=24 for all simulations.

## Discussion

The ability to respond to changes in social density is a crucial behavioral component across taxa, from bacteria to humans (34, 48–50), offering flexibility in coping with changing social environments. In zebrafish, young larvae swim in overdispersed groups, presumably to optimize their personal space and to avoid potentially harmful collisions with each other (12, 39). Nonetheless, the tendency to disperse and maximize nearest neighbor distance ought to be regulated to match different contexts. For example, tight aggregation might be unavoidable in the presence of local attractors such as food sources, or restricted areas of preferred temperature, luminance, flow or salinity. It is critical in such cases to attenuate and balance the drive to disperse with the tendency to aggregate around a spatially attractive location. Since the spatial profile of such sensory attractants usually changes slowly over time, the corresponding time constants with which desired population density setpoints are adjusted should change with comparable dynamics.

Interestingly, we find that the temporal modulations of inter-individual interactions of young larvae indeed have relatively long time constants, with notable modulations in behavior appearing after tens of minutes and lasting for several hours. We hypothesize that these long time constants are related to the natural occurrences of density modulation in the larvae’s environment, allowing fish to filter out and ignore fast, abrupt and stochastic changes in local population densities (51).

The mechanisms that allow zebrafish to estimate their population density are largely unknown. A few studies have looked into the role of olfactory (52) and somatosensory (38, 39) cues that larval zebrafish can use to estimate the presence of neighbors. In the current study, we show that fish also strongly rely on visual information to estimate population density by integrating looming events on their retina over time. In contrast, translational motion and retinal occupancy estimations, that were shown to guide the acute turning responses of fish to their neighbors (12, 41), seemed to play no role in modulating and adjusting the strength of repulsive interactions (Fig. 3). Notably, visual looming events are particularly informative in identifying conspecifics invading one’s ‘personal space’ since the invader’s perceived expansion velocity is a highly non-linear function of its distance (46). We found that modulations of avoidance behavior induced by such persistent looming events cause stable changes in collective behaviors. We predict that our findings will also generalize to older ages of larvae, when fish begin to aggregate and to exhibit both repulsive and attractive interactions with their neighbors, as early as 14 dpf (12, 13, 41). At these later developmental stages, exposure to high densities might cause fish to similarly attenuate only their repulsive interactions, resulting in tighter aggregations, or alternatively to attenuate both attractive and repulsive interactions, consequently forming more loose aggregations. Future studies can analyze history-dependent modulation in older zebrafish larvae and to test these different hypotheses.

To quantify the history dependent modulation of collective behavior we introduced a time varying state-space model (53, 54) that accurately describes both the static and dynamic features of collective behavior in varying social densities. This model deviates from the common approaches to modeling collective motion which rely only on static interaction rules (3, 5). Here, we propose a transition to a more general framework for describing collective motion, that can naturally incorporate additional modulating factors such hunger or stress, and varying environmental contexts such as presence of prey, water currents and changes in temperature.

This algorithmic modeling framework facilitates the generation of realistic circuit models that implement the underlying computations (12, 55). An explicit example of such a model, that implements directed turning in response to retinal occupancy, has been described previously (12). Here, we provide the experimental and behavioral framework that allows us to extend this particular circuit model to include how such turning behavior is modulated by exposure to persistent looming events. A core element of the extended model would be a population of looming sensitive neurons that exerts modulatory control on the critical, turning inducing nodes. Explicit circuit implementations underlying the detection of looming objects have been described previously in various organisms, including 7dpf larval zebrafish (43, 46, 47). We propose that the dedicated retino-tectal circuitry in the zebrafish is well poised to extract looming specific information (43, 47, 56). Our results suggest that looming events can then be integrated over longer time scales and lead to the updating of modulatory circuits possibly located in the dorsal raphe, the habenula or the hypothalamus. Such modulatory networks are well poised to change internal-state dependent output patterns as described, for example, for stress and hunger (57, 58).

The validation of such anatomically inspired circuit models requires functional interrogation at a brain wide level and at cellular resolution in a behaving animal (55, 59, 60). Our ability to detect experience dependent modulation of collective behaviors as early as 7 dpf, facilitates such investigation of the underlying biological and neural mechanisms, since larval zebrafish at this early stage are especially amenable to various brainwide neuroimaging and optogenetic perturbation approaches (61).

Our description of experience-dependent modulation of social interactions in larval zebrafish shows that even young animals, which are sometimes considered to have only a few simple and reflexive behaviors, can achieve increased behavioral diversity and cope with dynamic conditions. We hypothesize that other behavioral paradigms such as position holding in a moving stream and hunting moving prey might also be influenced by internal state variables such as we described here for population density (42, 62, 63). The behavioral assays and modeling framework developed here lend themselves readily to conduct future studies testing these hypotheses.

## Acknowledgments

We thank all members of the Engert and Fishman labs for support and advice during the project. We specifically thank Kumaresh Krishnan for his valuable help in establishing the behavioral setups. Finally, we would like to thank Ed Soucy, Brett Graham and Yuwei Li from the neuroengineering core facility at the Center for Brain Science at Harvard for their technical support. Roy Harpaz received funding from Harvard Minds Brain and Behavior initiative. Florian Engert received funding from the National Institutes of Health (U19NS104653 and 1R01NS124017-01), and the Simons Foundation (SCGB 542973 and NC-GB-CULM-00003241-02).

## Author Contributions

RH, MP, MCF and FE designed research, RH, MP and RG performed experiments, RH and FE analyzed the data, RH, MCF and FE wrote the paper.

## Supplementary figures

**Figure S1.**
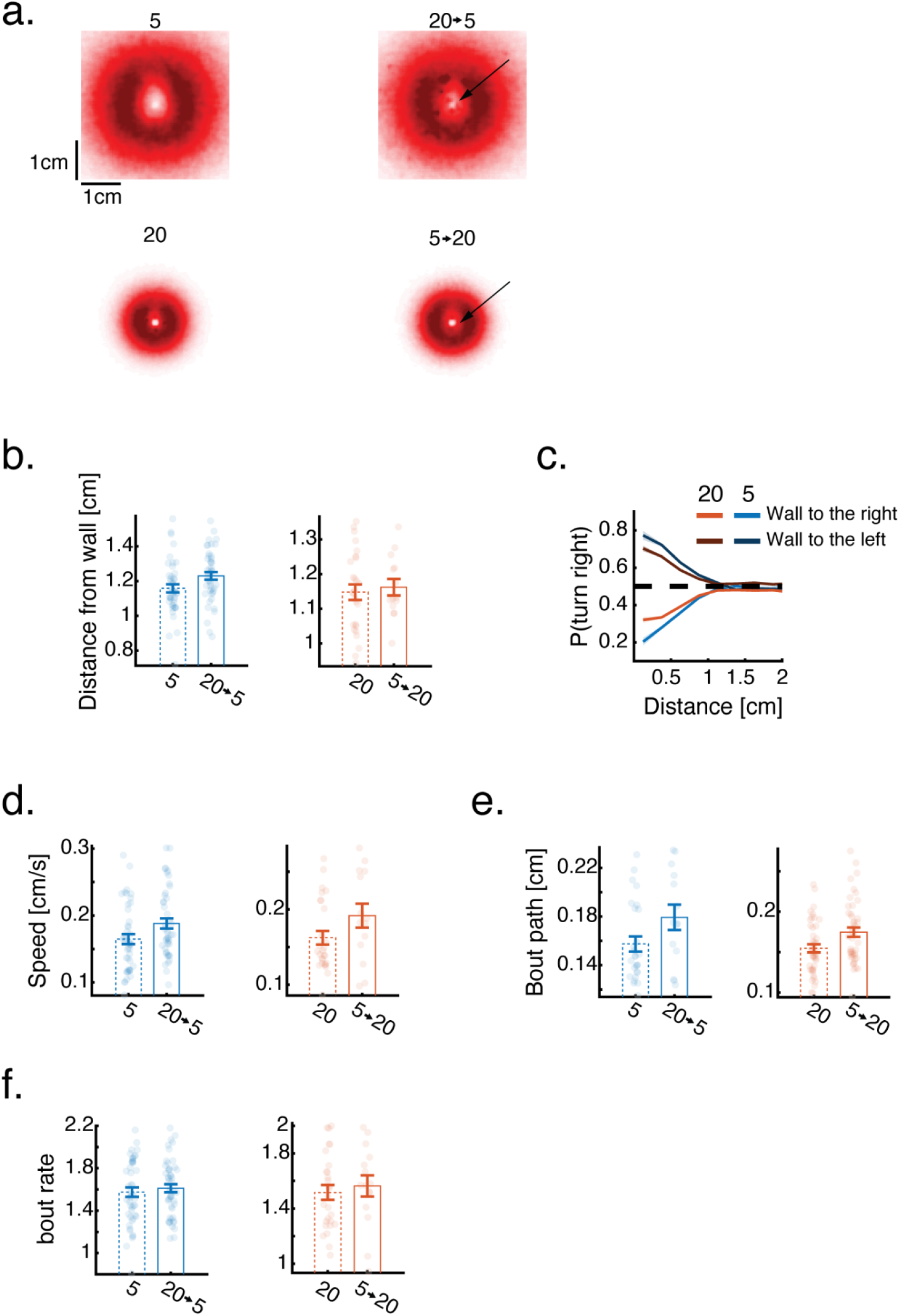
Previous social experience modulates collective behavior. **a.** Density maps depicting the 2d positions of nearest neighbor fish with respect to a focal fish situated at the center of the map pointing north. Maps are calculated from fish swimming in low (top row) and high (bottom row) neighbor densities that were either exposed to the opposite neighbor density in the past (right column) or to a similar density. Arrows point to areas of marked differences in density maps due to previous social experiences. **b.** Average wall distance of groups (Wall distance_5_=1.16±0.16, Wall distance_20→5_=1.23± 0.15 [mean±SD], p=0.0143, Wilcoxson’s rank sum test; Wall distance_20_=1.14±0.12, Wall distance_5→ 20_=1.16±0.09 [mean±SD], p=0.77, Wilcoxson’s rank sum test). **c.** Probability to turn right as a function of distance and direction (left or right) to the nearest wall for groups of 5 (blue colors) and 20 (red colors) fish. Lines represent mean turning probability calculated as the fraction of right turns out of all turns in 0.25 cm bins. Shaded areas are SEM. **d.** Average swimming speed of fish in groups (Speed_5_=0.16±0.05, Speed_20→5_=0.19±0.05 [mean±SD] p=0.0287, Wilcoxson’s rank sum test; Speed_20_=0.16±0.05, Speed_5→20_=0.19±0.06 [mean±SD], p=0.11, Wilcoxson’s rank sum test). **e.** Average path traveled in a bout of fish in groups (Bout path_5_=0.15±0.03, Bout path_20→5_=0.17±0.04 [mean±SD], p=0.0157 Wilcoxson’s rank sum test; Bout path_20_=0.16±0.03, Bout path_5→20_=0.18±0.03 [mean±SD], p=0.0962, Wilcoxson’s rank sum test) **f.** Average bout rate of fish in groups (Bout rate_5_=1.57±0.3, Bout rate_20→5_=1.61±0.26 [mean±SD], p=0.71 Wilcoxson’s rank sum test; Bout rate_20_=1.51±0.27, Bout rate_5→20_=1.56±0.29 [mean±SD], p=0.42, Wilcoxson’s rank sum test). In panels (b,d-f) Dots represent individual groups, error bars are mean±SEM.

**Figure S2.**
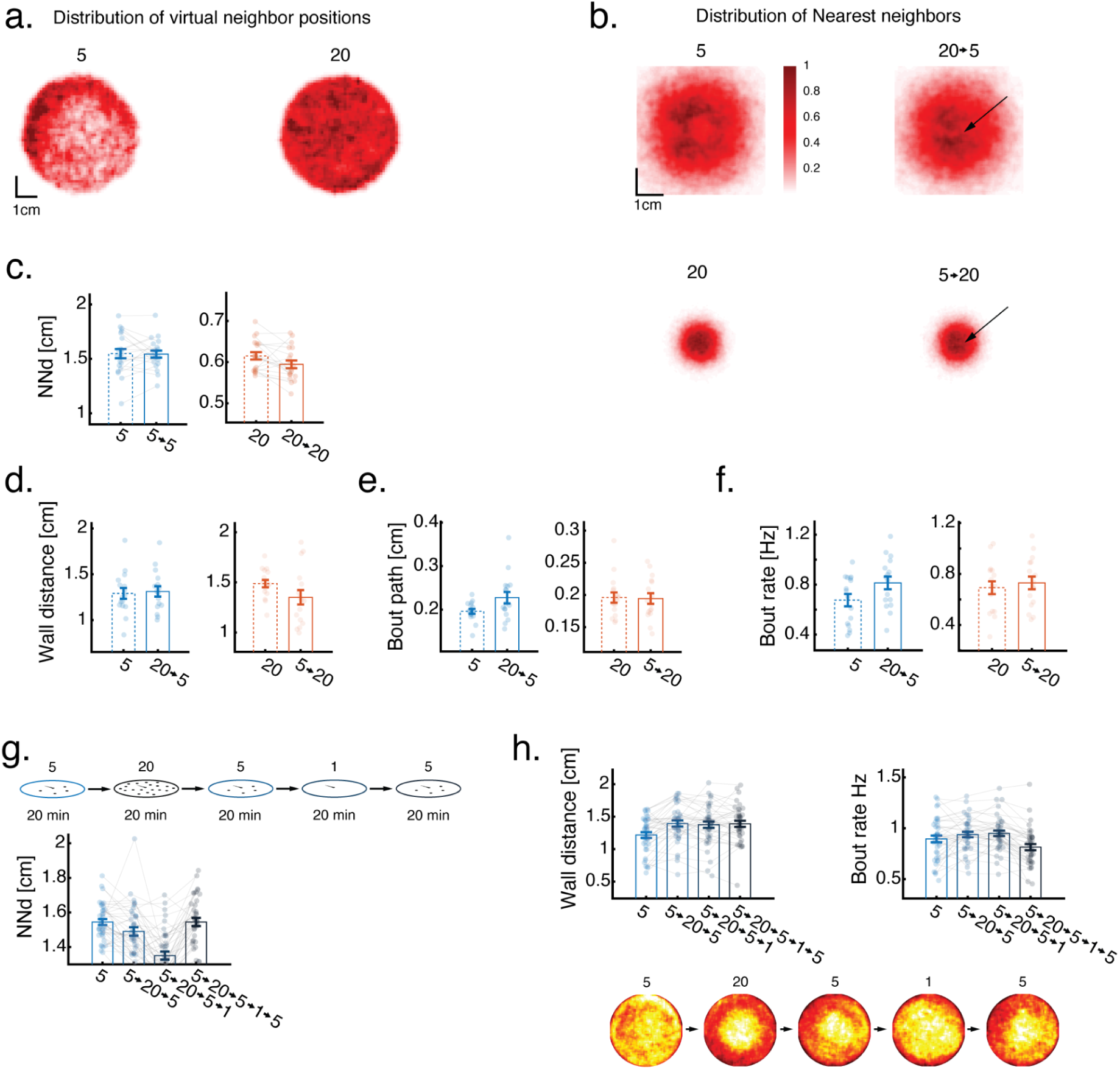
Virtual reality reveals the temporal characteristics of experience dependent modulation. **a.** Position density of real fish trajectories used as stimulus fish in VR experiments. Trajectories are characterized by some naturally occurring inhomogeneities. Using simulated trajectories instead of real fish trajectories, did not qualitatively change our findings. **b.** Density maps depicting the 2d positions of virtual nearest neighbors with respect to a focal fish situated at the center of the map pointing north. Maps are calculated from fish swimming in low (top row) and high (bottom row) virtual neighbor densities that were either exposed to the opposite neighbor density in the past (right column) or to a similar density (left column). Arrows point to areas of marked differences in density maps due to previous social experiences. **c.** Average nearest neighbor distance of real fish that did not experience a change in virtual neighbor density (NN_5_=1.547±0.19, NN_5→5_=1.544±0.14 [mean± SD], p=1 Wilcoxson’s signed rank; NN_20_=0.615±0.04, NN_20→20_=0.594±0.04 [mean±SD], p=0.02 Wilcoxson’s signed rank). **d.** Average wall distance of real fish (Wall distance_5_=1.29±0.23, Wall distance_20→5_=1.31±0.22 [mean±SD], p=0.9 Wilcoxson’s rank sum; Wall distance_20_=1.49±0.15, Wall distance_5→20_=1.35±0.29 [mean±SD], p=0.057 Wilcoxson’s rank sum). **e.** Average path traveled within a bout of real fish (Bout path_5_=0.2±0.02, NN_20→5_=0.22±0.05 [mean±SD], p=0.067 Wilcoxson’s rank sum; Bout path_20_=0.2±0.03, NN_5→20_=0.2±0.03 [mean±SD], p=1 Wilcoxson’s rank sum). **f.** Average bout rate of real fish (Bout rate_5_=0.67±0.2, NN_20→5_=0.82±0.2 [mean±SD], p=0.062 Wilcoxson’s rank sum; Bout rate_20_=0.7±0.2, NN_5→20_=0.73±0.2 [mean±SD], p=0.777 Wilcoxson’s rank sum). **g.** Nearest neighbor distance of real fish before and after exposure to high density and after subsequent exposure to 0 virtual neighbors (NN_5_=1.54±0.1, NN_5→20→5_=1.49±0.15 [mean±SD], p=0.005 Wilcoxson’s signed rank; NN_5_=1.54±0.1, NN_5→20→5→0→5_=1.54±0.15 [mean±SD], p=0.7 Wilcoxson’s signed rank). **h. Top:** Wall distance, bout rate and position density of real fish before and after exposure to high density and after subsequent exposure to 0 virtual neighbors (Wall distance_5_=1.22±0.28, Wall distance_5→20→5_=1.39±0.29 [mean±SD], p=0.0002 Wilcoxson’s signed rank; Wall distance_5→20→5_=1.39±0.29, Wall distance_5→20→5→0→ 5_=1.39±0.29 [mean±SD], p=0.81 Wilcoxson’s signed rank). **Bottom:** position density maps of real fish in the 5 subsequent experimental conditions. In panels (c-h) dots represent fish, lines connect data from same fish, error bars are mean±SEM.

**Figure S3.**
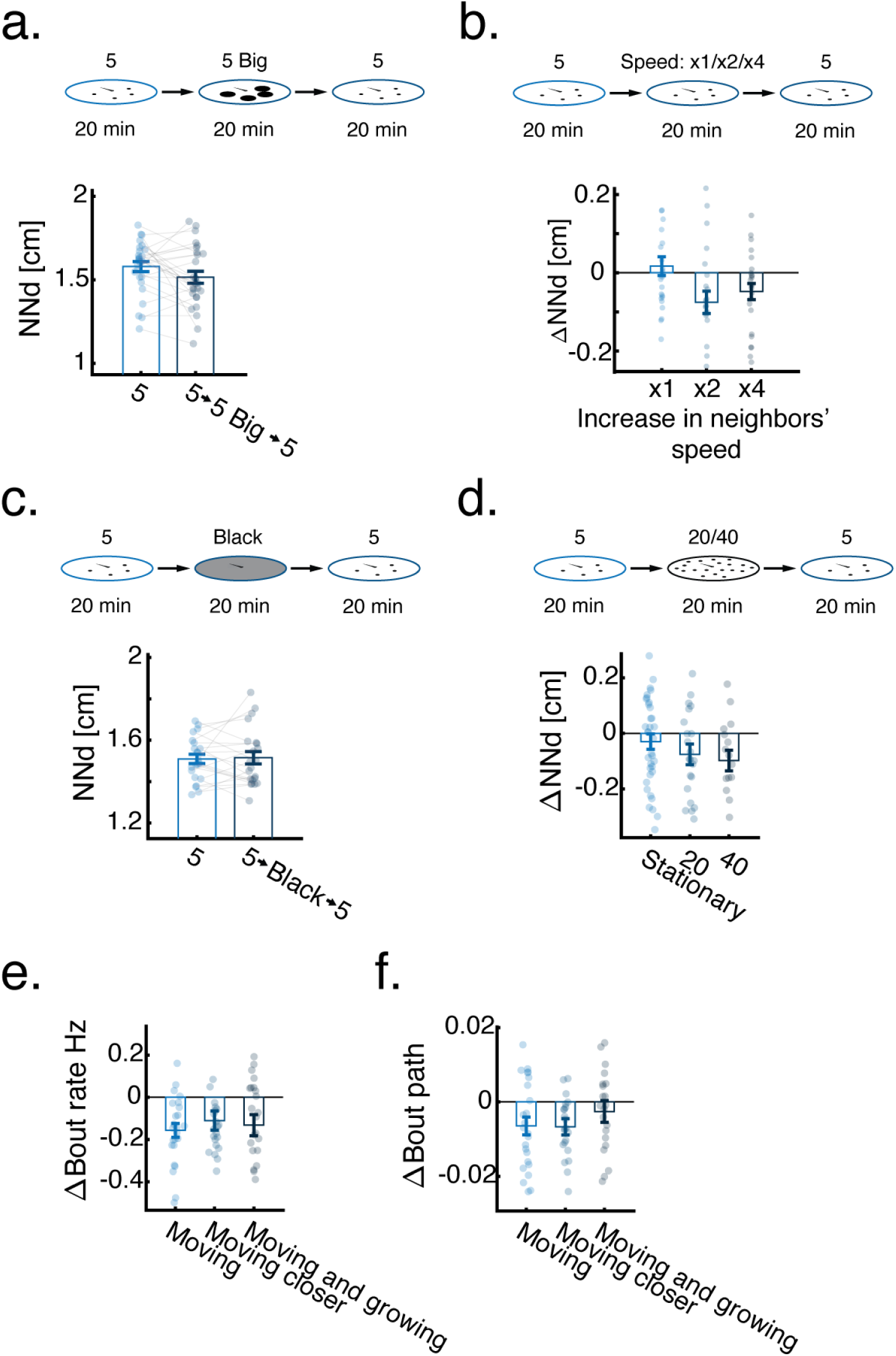
Persistent changes in retinal occupancy drive experience-dependent modulation of collective behavior. **a.** Average nearest neighbor distance of fish swimming in a low density group before and after 20 min of exposure to the same number of virtual neighbors, but with a larger size - r_large_=4·r_small_ (NN_5_=1.58±0.15, NN_5→5 big→5_=1.51±0.18 [mean±SD], p = 0.067 Wilcoxon signed rank). **b.** Average change (after-before) in nearest neighbor distance of fish swimming in a low density group after 20 min of exposure to the same density of neighbors swimming in different speeds (ΔNN_speed x1_=0.017±0.12, ΔNN_speed x2_=-0.076±0.0.14, ΔNN_speed x4_=-0.048±0.1 [mean±SD], p_speed x1_=0.8, p_speed x2_=0.013, p_speed x4_=0.032 Wilcoxson signed rank). **c.** Average nearest neighbor distance of fish swimming in a low density group before and after exposure to 20 min of a black whole-field background (NN_5_=1.51±0.1, NN_5→black→5_=1.515±0.14 [mean±SD], p = 0.98 Wilcoxon signed rank). **d.** Average change (after-before) in nearest neighbor distance of fish swimming in a low density group after exposure to 20 min of different high densities of neighbors (ΔNN_stationary 20_=0.032±0.13, ΔNN_20_=-0.075± 0.18, ΔNN_40_=-0.1±0.15 [mean±SD], p_stationary 20_=0.387, p_20_=0.059, p_40_=0.03 Wilcoxson signed rank). **e.** Average change (after-before) in bout rate of fish swimming in a low density group after exposure to 20 min of high density of neighbors with different motion statistics (Δbout rate_moving_=-0.16±0.16, Δbout rate_moving closer_=-0.11±0.22, Δbout rate_moving and growing_=-0.13±0.24 [mean±SD], p_moving_=0.0003, p_moving closer_=0.0072, p_moving and growing_=0.015 Wilcoxson signed rank). **f.** Average change (after-before) in bout path of fish swimming in a low density group after exposure to 20 min of high density of neighbors with different motion statistics (Δbout path_moving_=-0.0064±0.012, Δbout path_moving closer_=-0.0067±0.011, Δbout path_moving and growing_=-0.0026±0.014 [mean±SD], p_moving_=0.018, p_moving closer_=0.0033, p_moving and growing_=0.465 Wilcoxson signed rank). In panels (a-f) dots represent fish, error bars are mean±SEM.

**Figure S4.**
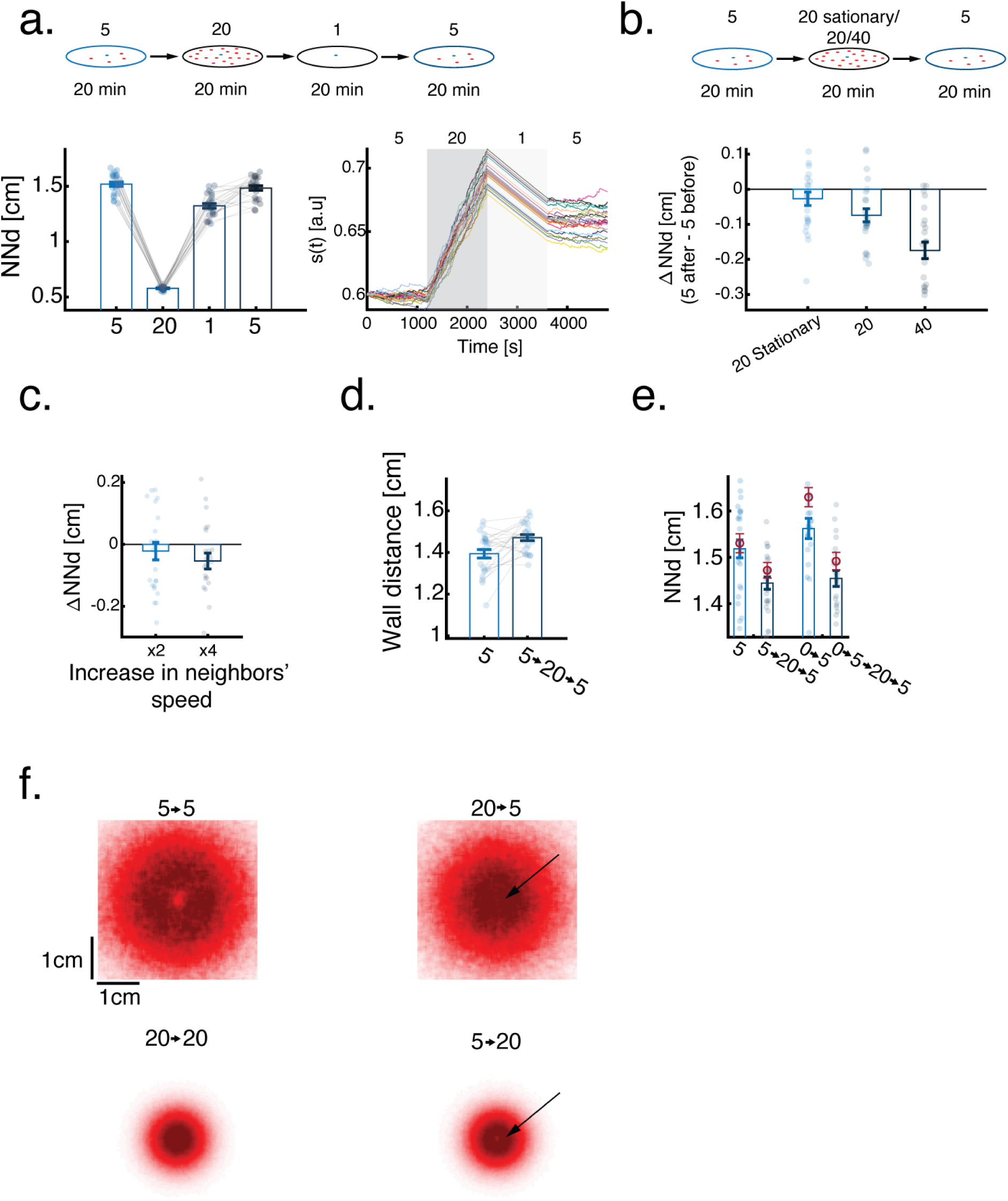
Models accurately predict experience dependent modulation of collective behavior. **a. Left**: Nearest neighbor distances of simulated agents swimming in different densities of virtual neighbors starting from low (4 neighbors), switching to high (19 neighbors), 0 neighbors and returning to low again. Dots represent different simulated agents (N=24). **Right:** Value of the internal state variable *s*(*t*) over time for the same agents shown on the left. Different colors represent different simulated agents responding to the same neighbor stimuli. *s*(*t*) increases sharply with the switch to high density and decays more slowly with the switch to 0 neighbors. After switching back to low density it remains stable for the 20 minutes period. **b.** Average change (after-before) in nearest neighbor distance of fish swimming in a low density group after exposure to 20 min of different high densities of neighbors. **c.** Average change (after-before) in nearest neighbor distance of agents swimming in a low density group after exposure to a low density of virtual neighbors swimming at different speeds. **d.** Average wall distance of agents swimming in a low density group before and after exposure to a high density of virtual neighbors. **e.** Average nearest neighbor distance of agents swimming in a low density group before (light blue) and after (dark blue) exposure to a high density of virtual neighbors, and with (right) or without (left) pre-exposure to 0 neighbors. Dark red circles and error bars represent mean± SEM of the real fish data from similar experiments. **f.** Density maps depicting the 2d positions of nearest neighbors with respect to a focal simulated agent situated at the center of the map pointing north. Maps are calculated for agents swimming in low (top row) and high (bottom row) agent densities that were either exposed to the opposite neighbor density in the past (right column) or to a similar density (left column). Arrows point to areas of marked differences in density maps due to past experience. In panels (b-e) dots represent simulated fish, error bars are mean±SEM and N=24 simulation repetitions.

## Methods

### Fish husbandry

All larvae used in the experiments were obtained by crossing adult AB zebrafish. Larvae were raised in low densities of approximately 40–50 fish in large petri dishes (D = 12 cm). Dishes were filled with filtered fish facility water and were kept at 28°C, on a 14–10h light dark cycle. From age 5 dpf, fish were fed paramecia once a day. All experiments followed institutional IACUC protocols as determined by the Harvard University Faculty of Arts and Sciences standing committee on the use of animals in research and teaching.

### Group swimming experiments with density switches

At age 7dpf, fish were transferred from their holding dishes to round custom-designed experimental arenas (d = 6.5cm, depth 1cm), filled with filtered fish facility water up to a height of ∼0.8 cm (12). Briefly, these plastic experimental arenas had a flat bottom and curved walls (half a sphere of radius 0.5 cm) and were sandblasted to prevent reflections and allow for stimulus projection directly on the arena’s surface (see (12) for full details). Every experimental arena was filmed using an overhead camera (Grasshopper3-NIR, FLIR System, Zoom 7000, 18–108 mm lens, Navitar) and a long pass filter (R72, Hoya). Arenas were lit from below using 4 infrared LED panels (940 nm panels, Cop Security) and from above by indirect light coming from 4 32W fluorescent lights. All recording procedures and subsequent fish tracking were performed as described in (12).

Fish were pre-acclimated in either low density (5 fish) or high density (20 fish) for 4 hrs prior to behavioral imaging. After the pre-acclimation period, fish were imaged for 20 minutes to obtain a baseline measure of their behavior. We then combined 5 fish groups into 20 fish groups or separated 20 fish groups into 5 fish groups. Fish were then imaged again for 20 minutes to measure how pre-exposure to one density affects behavior in the current density. Groups were eliminated from subsequent analysis in the case that one or more of the fish were immobile for more than 50% of the experiment. All and all 7.2% and 9.3% of groups of 5 and 20 fish were eliminated from the analysis due to immobility of the fish. Choosing a more stringent, or a less stringent criteria for elimination did not change the qualitative nature of the results.

### Fish kinematics and group level properties

Individual and group level properties were extracted following the procedures described in (12). The position of each fish *i* at time *t* is defined as the center of mass of the fish extracted from offline tracking and is denoted as 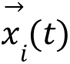. The velocity of each fish *i* is given by 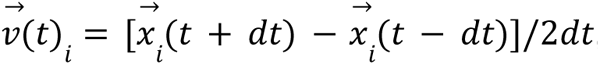, where dt is 1 frame or 0.033s. The speed of the fish is then 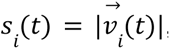, and the direction of motion is 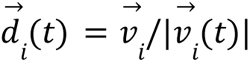. Distance between fish *i* and *j* is the euclidean distance 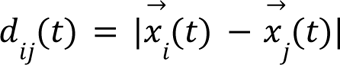. Chance levels for interindividual distances were calculated from randomized groups created by mixing fish from separately recorded groups and randomly shuffling their time signatures. For 20 fish groups, we allocated fish from different groups to the extent possible by the number of tested groups, filling in with fish that did swim together but randomly shuffling their time signatures.

### Estimating retinal occupancy

We defined the effective retinal field of each eye as the visual angle extending from the fish’s heading direction (0°) towards its tail (±165°), assuming a 30° blind angle directly behind the fish. To estimate retinal occupancy, we used the physical length of a neighboring fish (a straight line from snout to tail), the distance to the focal fish and the relative angle with respect to the heading direction of the focal fish to calculate the specific angles out of the retinal field occupied on each eye. When estimating total occupancy within each eye we corrected for instances of neighboring fish occluding one another (Fig. 1e).

### Turning in response to the difference in retinal occupancy between the eyes

To estimate turning direction (left vs. right) in response to difference in retinal occupancy between the eyes P(turn right| Δretinal occupancy)(Fig. 1e) we measured the change in heading angle 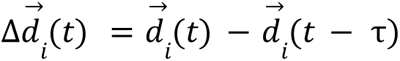 around a detected bout event and used the visual occupancy measured at the beginning of that bout. Specifically, P(turn right| Δretinal occupancy) is the fraction of right turns out of all left/right turns recorded for 5° (groups of 5 fish) or 10° (groups of 20 fish) bins of Δretinal occupancy. We discarded all turning events at distance < 1.25 body length from the wall, as not to confound wall avoidance with neighbor responses.

### Virtual reality experiments

Every VR experimental arena was filmed using an overhead camera (Grasshopper3-NIR, FLIR System, Di II VC, 18–200 mm lens, Tamron) and a long pass filter (IR720, Zuma). All experimental arenas were lit from below using an infrared LED panel (850 nm panels, Univivi) and from above by indirect light coming from 4 32W fluorescent lights. Stimuli were projected from below, directly onto the sandblasted arenas using P7 mini LED projectors (AAXA) (Fig. 2a). Every 8 experimental arenas were connected to a single computer controlling stimulus projection, video recording and online behavioral tracking. Stimuli were updated at 60hz and behavioral data was recorded at 90hz. All and all we used 24 experimental arenas allowing us to test different experimental conditions on the same experimental day within the same clutch of fish. All fish tracking and posture analysis were done using custom software written in Python 3.7 and OpenCV 4.1 as described in (12, 55). Briefly, movie images were background subtracted online to obtain an image of the swimming fish, and body orientation was estimated using second-order image moments. In all experiments, fish were first pre-acclimated in their designated groups (5 or 20 real fish) for at least 4 hrs. Single fish were then transferred to individual arenas and interacted with the projected virtual neighbors. Fish with bout rates < 0.25Hz were removed from the analysis, as 7dpf larvae are expected to bout at around 1Hz. All and all less than 5% of the fish were discarded from our analysis. To filter out instances where fish were temporarily immobile, we limit our data to instances when fish actively performed bouts. While this filtering removes some noise in our data it does not change any of our findings.

### Trajectory replay stimuli

Virtual neighbors were presented as dark dots (d=1.5cm) on a light background. We used example trajectories of 5 and 20 fish, randomly chosen from real groups recorded in the free swimming experiments. In cases where stimuli presentation time in the experiments exceeded the length of the recorded trajectories (20min), we restarted the stimulus causing a momentary discontinuity in the presented stimuli. Similarly, when switching between low and high densities, a momentary discontinuity exists in the dots’ trajectories. To confirm that our findings are not unique to the specifically chosen stimuli sets, we created surrogate fish trajectories by randomly sampling from the distribution of bout sizes, interbout intervals and turning angles of the recorded fish. Experiments using these surrogate data sets, closely matched our findings using the replayed trajectories. Below we describe the various stimuli sets we used in all VR experiments:

- *Increasing time of high density* (Fig. 2e). We repeatedly presented a 5 min segment of the 20 min trajectories to better compare between shorter and longer exposures. For example, fish that were exposed to 20 min of high density, were presented with the same 5 min stimuli as fish that were exposed to 5 min of high density, but the stimuli looped 4 times for the 20 min condition.
- *Stationary dots* (Fig. 3a, Movie 3). We used a single, randomly chosen frame from the recorded fish trajectories. Dots were stationary for the entire stimulus presentation time.
- *Low density with increased speed* (Fig. S3b). To speed up fish swimming by a factor of 2 or 4, we downsampled the recorded trajectories. This allowed fish to traverse the same distance by either 1/2 or 1/4 of the time. We then looped the trajectories to compensate for the shorter total presentation time.
- *Rotational trajectories* (Fig. 3a, Movie 3). To independently test the effects of rotational energy we presented a high density of dots (19) moving on 1d rings centered around the focal fish. Radial distances from the focal fish were chosen based on the most likely distance to find the 1st, 2nd…19th nearest neighbor in the recorded trajectories. Angular position with respect to the heading direction of the focal fish was changed in a bout-like fashion, with clockwise/counter clockwise direction of motion chosen randomly for each bout.
- *Stationary Looming dots* (Fig. 3a, Movie 3). To independently test the effects of looming energy, we arranged 19 dots equally spaced from one another within a circle (forming an hexagonal formation). Dot sizes increased and decreased with temporal dynamics taken from the recorded changes in angular size experienced by a focal fish swimming with virtual neighbors. Minimum and maximum dot radii were chosen at *r_max_* = 0. 9cm and *r_min_* = 0. 075cm.
- *Trajectories of 40 agents* (Fig. S3d). We combined the real recorded trajectories of two 20 fish groups to form a 40 agent group.

### Single dot stimulus

We used a similar protocol for single dot presentations as described in (12). Briefly, In each trial, a single dot appeared at one side of the fish at a given radial distance and moved with fish-like bouts tangentially around the fish from ±75° to ±30° for 5s, with 0° taken as the fish heading direction. Presentation side was chosen at random and once a dot reached the end of its trajectory a new dot immediately appeared (0s inter-stimulus interval) (Fig. 3b, Movie 4). We manipulated the dot’s radial distance or the dot’s radius according to the following stimulus conditions:

● *‘Moving’ dot stimulus.* We set dot radius = 0.15cm and distance to dot center = 0.42cm. Hence, dots had a constant angular size of ∼40° throughout the trial.
● *‘Moving closer’ dot stimulus.* Dots had a constant radius = 0.075cm, and the distance to the dot center gradually decreased over the 5s presentation time from 0.92cm to 0.22cm, i.e. angular size changed from ∼9° to ∼40° throughout the trial.
● *‘Moving and growing’ dot stimulus.* Dots were presented at a constant distance of 0.35 cm, and dot radius was increased from 0.025cm to 0.13, i.e. angular size changed from ∼9° to ∼40° throughout the trial.

### Estimating fish response to a single dot stimulus

To analyze fish responses to the single dot stimuli we calculated for each bout during stimulus presentation the change in body orientation of the fish. We then calculated the overall probability to turn left or right in a given trial. These probabilities were then averaged over blocks of trials and over fish (Fig. 3b).

### Estimating fish responses to global retinal occupancy based on monocular averaging and binocular comparisons

Previously, we showed that 7 dpf larvae turn away from the eye that experiences a higher retinal occupancy (12). Responses to the global retinal occupancy of neighbors occupying the set of visual angles θ^*left*^, θ^*right*^ (on the left and right eyes) follow the form:

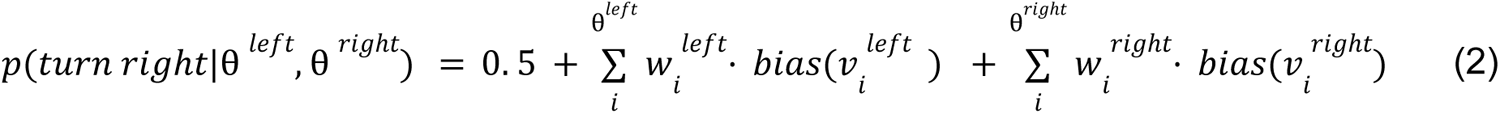

Where 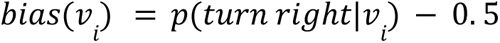 is the turning direction bias in response to vertical retinal occupancy *v_i_*, *w_i_* is the relative weight assigned to each response bias and represents a weighted average 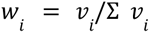 of the turning biases for each eye. The intercept 0.5, centers the summed responses around that value and 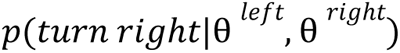, is bounded between 0 and 1. If we assume that 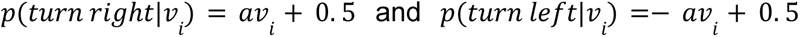, i.e. response can be approximated as a linear function of *v_i_*, we can rewrite the above equation as:

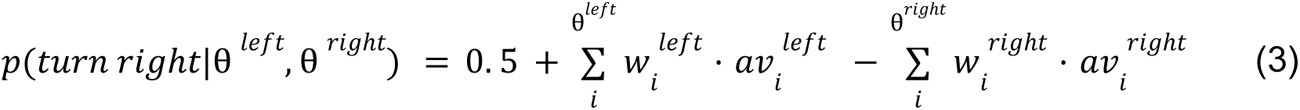

And since 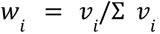, we get:

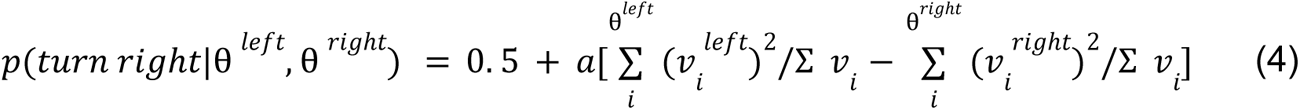

Hence, the turning direction of the fish is dependent linearly on the difference in the weighted average of the squared vertical visual angles in each eye. We estimate this function from data in Fig. 4b and use it in simulating fish responses to virtual neighbors (see below) (Fig. 5a-d).

### Time varying models of fish social interactions

The basic individual agent swim properties, responses to arena walls and static inter-individual interactions based on retinal occupancy used in the models here, were previously described in (12). We briefly describe the details again here, and expand on the novel aspects of experience dependent modulation of inter-individual interactions.

a. *Basic swim properties*. Each stationary fish, at every time step, probabilistically decides to perform a bout according to the average bout rates of real fish. Similarly, bout duration and distance traveled in a bout followed that of the average bouts of real fish (12).
b. *Wall interactions*. Similar to real fish, simulated agents’ first priority is to avoid crushing into the walls. Therefore, agents ignore their neighbors when in close proximity to the arena walls and turn away from the direction of the closest arena wall. In every bout, the probability to respond to the walls and to turn in the opposite direction is drawn from the empirical wall response functions observed in real fish (Fig. S1c). If the executed bout was expected to end outside of the arena, it was truncated to ensure the fish stayed inside the simulated arena.
c. *Turning behavior in response to global retinal occupancy of neighbors*. We used the algorithms described in (12) and in Eq. 2 (see above) to simulate the response of agents to the global retinal occupancy of their neighbors. When modeling agents responding to stimuli from VR experiments (Fig. 5a-d), we used the estimated response to vertical retinal occupancy extracted from VR experiments (Fig. 4a) and the linear approximation presented in Eq. 4. In models simulating a group of interacting agents (Fig. 5e-f), we used the estimated responses to vertical retinal occupancy described in (12). Vertical retinal occupancy of neighbor i at visual sub-angle j (Vc_ji_) was calculated using the simulated height (H_j_) and distance (d_j_) of neighbor 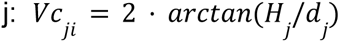 . For simplicity, we did not account for occlusions among neighbors in estimating retinal occupancy as initial simulations showed that it did change our simulation results. In addition, we also assume that the height of the fish along its body axis is constant, allowing us to treat the vertical occupancy at all visual sub-angles occupied by neighbor j as a single value.
d. *Leaky Integrator module for temporal integration of neighbor-evoked looming events*. The experimental findings from VR experiments (Fig. 2e-g, Fig. 3a-c) indicate that agents slowly integrate (and remember) neighbor-evoked looming events on their retina. This process develops slowly, remains stable for many hours, and can be reverted back to baseline via the removal of all retinal inputs (Fig. 2e-g). We therefore introduce a leaky integrator module, that temporally updates an internal state variable *s*(*t*) according to the the input to the integrator *input*(*t*), which is the maximal increase in retinal occupancy of any neighbor experienced at time *t* (Eq. 1). In the absence of visual input (or with a weak input), the internal state variable will exponentially decay back to a resting state value *s_rest_* with a time constant τ.
e. *Linking internal state s*(*t*) *to agents’ occupancy based turning behavior*. We assume that higher internal state values *s*(*t*) indicate weaker responses to retinal occupancy of neighbors, as seen in behavioral experiments. Therefore we linked *s*(*t*) to the strength of response to retinal occupancy 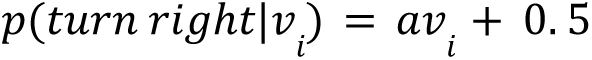. For simplicity, we assumes a linear relationship between *s*(*t*) and the slope of the response *a*, and define it as 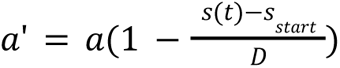 is the starting value of the integrator, D is a scaling factor and α is the value estimated from groups of 5 fish that were not previously exposed to changes in social density (Fig. 4b)(12).
f. *Switching simulated agents between high and low density*. When switching a group of simulated agents between densities mid simulation (Fig. 5e-f), we either removed (high to low) or increased the number of simulated agents. When removing agents, we simply chose *N_to remove_* agents randomly, and eliminated them from the rest of the simulations. When adding agents, we assigned the *N_to add_* agents starting positions and heading orientations at random. The internal state values *s*(*t*) of the added agents, were chosen such that they represent the distribution of values of internal states of the agents already swimming together. Therefore, we randomly drawn *s*(*t*) for the newly added agents from a normal distribution with mean and sd calculated from *N* agents already swimming together.
g. *Model parameters used in simulations*

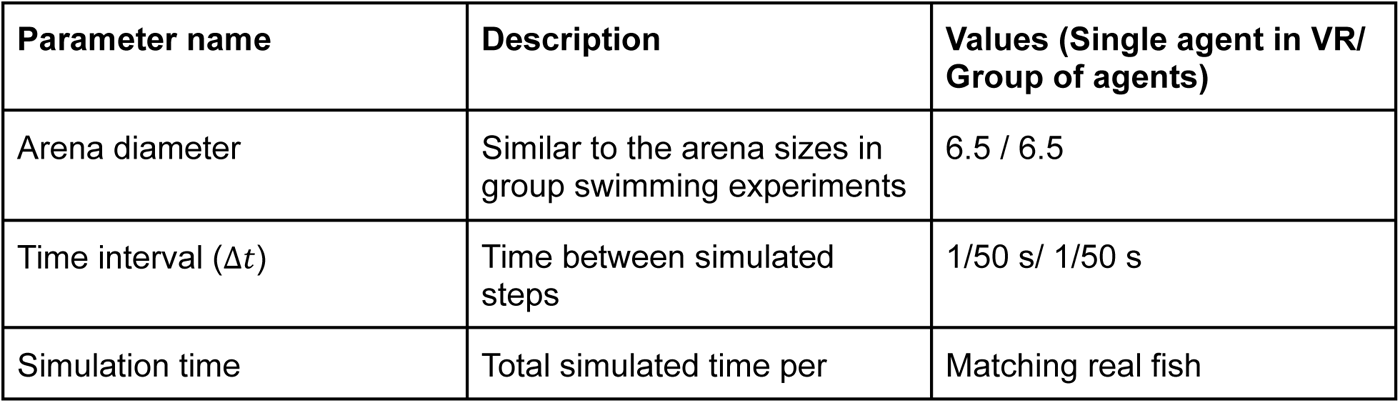
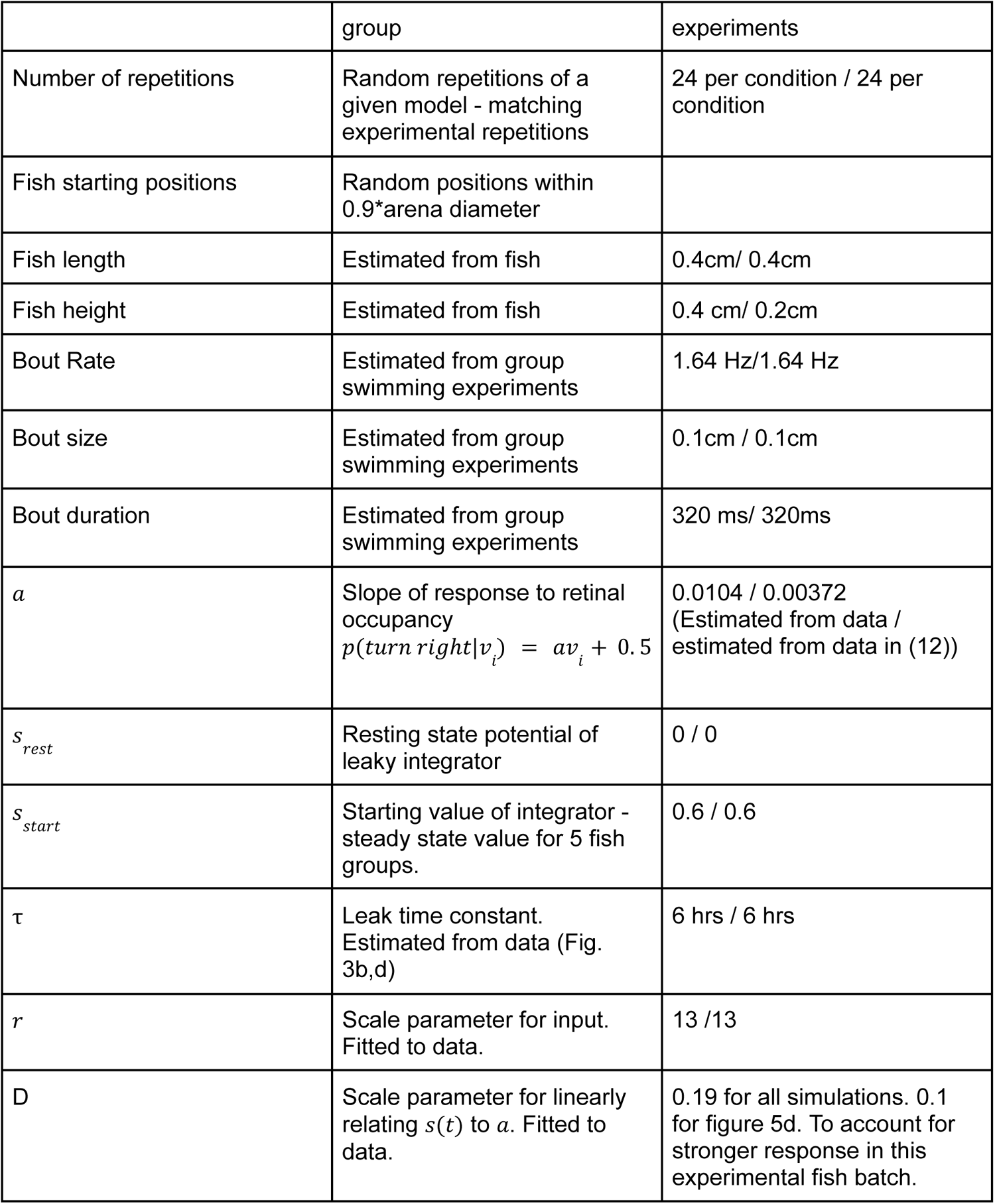

### Sample sizes, and power estimation

Sample sizes for group swimming experiments were chosen to allow accurate estimates of group level statistics (e.g. nearest neighbor distances) and inter-individual interaction functions according to previously reported data in larval zebrafish (12, 41), and to allow at least 10 degrees of freedom when using non-parametric statistical models to compare between experimental conditions. Due to the unique experimental structure (where 5 fish groups are combined to 20 fish groups), there is considerably more data collected for 5 fish groups than 20 fish groups. In virtual reality experiments, we used 16-24 fish per experiment, in accordance with previously published data (12) and as our preliminary data showed that these numbers are sufficient to estimate fish responses and the differences between experimental conditions.

### Statistical testing

We used non-parametric statistical modeling to compare between experimental conditions and report exact p-values. We also report mean±sem, and plot all individual data points. We report Cohen’s d as a measure of effect size when appropriate. Before a statistical model was chosen we made sure model assumptions are fulfilled. All reported p-values, when no a-priori hypothesis of the direction of the effect exists, are two-sided p-values. Our treatment of statistical reporting followed the guidelines in (64).

## Code and data availability

All data are publicly available at: https://doi.org/10.7910/DVN/KWA7DK

All codes used for model simulations are publicly available at: https://github.com/harpazone/Experience-dependent-modulation-of-collective-behavior

